# Diversification, Spread, and Admixture of Octoploid Strawberry in the Western Hemisphere

**DOI:** 10.1101/2021.03.08.434492

**Authors:** Kevin A. Bird, Michael A. Hardigan, Aaron P. Ragsdale, Steven J. Knapp, Robert VanBuren, Patrick P. Edger

## Abstract

**Premise of the study:** Octoploid strawberry (*Fragaria sp*.) has a complex evolutionary history that has until recently been intractable due to limitations of available genomic resources. While recent work has further uncovered the evolutionary history of the octoploid strawberry, there are still open questions. Much is still unknown about the evolutionary relationship of the wild octoploid species, *Fragaria virginiana* and *Fragaria chiloensis*, and gene flow within and among species after the original formation of the octoploid genome.

**Methods:** We leveraged a diversity collection of wild octoploid ecotypes of strawberry representing the recognized subspecies and ranging from Alaska to Southern Chile, and a high density SNP array to investigate wild octoploid strawberry evolution. Evolutionary relationships are interrogated with phylogenetic analysis and genetic clustering algorithms. Additionally, admixture among and within species is assessed with model-based and tree-based approaches.

**Key Results:** Phylogenetic analysis revealed that the two octoploid strawberry species are monophyletic sister lineages. The genetic clustering results show substructure between North Americana and South American *F. chiloensis* populations. Additionally, model-based and tree-based methods support gene flow within and among the two octoploid species, including newly identified admixture in the Hawaiian *F. chiloensis* subsp. *sandwicensis* population that appears to be from an ancestral *F. chiloensis* population.

**Conclusion:** *F. virginiana* and *F. chiloensis* are supported as monophyletic and sister lineages. All but one of the subspecies recognized within both octoploid species show extensive paraphyly. Furthermore, the phylogenetic relationship among *F. chiloensis* populations supports a single population range expansion southward from North America. The inter- and intraspecific relationships of octoploid strawberry are complex and suggest substantial and deep gene flow between sympatric populations among and within species.

## Introduction

The ploidy of strawberrry, *Fragaria sp*., ranges from diploid to decaploid, and recent work has shown an impressive retention of karyotype and genome structure over tens of millions of years (Hardigan et al., 2020a). The prevalence of neopolyploids makes the genus a powerful system to study the immediate effects of polyploidy like subgenome dominance (Edger et al., 2019) and adaptability to new environments (Wei et al., 2019). Additionally, the ability to easily hybridize or duplicate genomes allows for experimental manipulation of ploidy level for ecological studies (Wei et al., 2020). The cultivated garden strawberry *Fragaria* × *ananassa* is unusual for a domesticated crop in that it is a homoploid hybrid of two wild octoploids and has a very recent domestication history (<300 years) that is well documented, so the progenitor species and populations are well known (Darrow, 1966; Pincot et al., 2021; Hardigan et al., 2020b).

*F. virginiana* and *F. chiloensis*, the wild octoploid progenitors of *F. × ananassa*, are native to the Western hemisphere. *F. virginiana* is distributed across North America, whereas *F. chiloensis* is only distributed along the coast of Western North America from Alaska to Southern California as well as Hawaii and the Chilean coast in South America (Staudt, 1988, 1999, 2008). It was these two species, *Fragaria virgniana* from North America and *Fragaria chiloensis* from South America, that would result in the spontaneous hybrid formation of the cultivated strawberry *F. × ananassa* throughout Europe in the 18th century after being transported from the Western hemisphere (Darrow, 1966; Pincot et al., 2021). Their wide range, including sympatry in North America where natural hybrids have been previously observed (Dillenberger et al., 2018), and role as progenitors to an important agricultural crop has produced great interest from evolutionary biologists and ecologists as well as plant breeders. However, because of the complex nature of the octoploid *Fragaria* genome, there has been limited investigation of these wild octoploids at the genetic and genomic level (Hardigan et al., 2020b). Despite the rich historical knowledge of these species, their well documented range and the observation of natural hybrids, there are many outstanding questions about the relationship and origins of these octoploid species and the nature of intra- and inter-specific gene flow between populations that can only be explored through genetic and genomic techniques.

Previous analyses have provided glimpses into the nature of the evolutionary relationships of these species. Dillenberger et al. (2018) assessed the phylogeny of several *Fragaria* species of different ploidy using whole plastomes. Their results suggest that *Fragaria virginiana* is poly- and paraphyletic and that *F. chiloensis* is monophyletic and derives from a *F. virginiana* subspecies. However, plastome phylogenies can differ from a true species tree under complex evolutionary scenarios because plastomes are uniparentally inherited and represent only a single marker. In recognition of this Dillenberger et al. (2018) note that hybridization, incomplete lineage sorting, or both may explain their observed phylogenetic relationships. Therefore, analysis of these populations using nuclear DNA is needed to clarify the phylogeny of these taxa and identify the presence and extent of gene flow. The recently published genome of *Fragaria × ananassa* and accompanying resources has allowed for previously intractable questions about the ancient diploid progenitors of the octoploid strawberry genome (Edger et al., 2019) and the domestication history of cultivated strawberry (Hardigan et al., 2020b) to be dissected. Hardigan et al. (2020b) used the program TreeMix to provide evidence of admixture between *F. chiloensis* from North American and *F. virginiana*, however more in depth investigation of gene flow within and among species and their phylogenetic relationships is still needed. There are many questions remaining about the relationships and evolution of these wild octoploids. It is unclear whether the two octoploid species are sister lineages or whether *F. chiloensis* derives from *F. virgniana* subspecies. A more detailed look at intra- and interspecific gene flow between octoploid subspecies, rather than broad geographic groupings, is needed to characterize movements and mixtures of these populations. Additionally, the *F. chiloensis* subsp. *sandwicensis* has not been analyzed thus far in any phlogenetic or population genetic analyses.

Here we leverage a recently published *F. × ananassa* genome (Edger et al., 2019) and 50K Axiom SNP array (Hardigan et al., 2020a) to study a phylogenetically diverse sample of *F. virginiana* and *F. chiloensis* populations collected throughout the natural geographic ranges of the underlying subspecies in North and South America. Using a variety of genetic and genomic methods, we show evidence that *F. virginiana* and *F. chiloensis* are monophyletic sister lineages, but subspecies designations show substantial paraphyly. We also demonstrate the extent of intra-specific gene flow between geographically diverged *Fragaria* populations. Notably, we provide novel evidence that the Hawaiian *F. chiloensis* subsp. *sandwicensis* experienced gene flow from ancestral *F. chiloensis* populations. These results build upon Dillenberger et al. (2018)’s previous work by supporting the monophyly of *F. chiloensis* while clarifying that *F. virginiana* is also monophyletic and that these are sister species. We additionally show the nature and extent of intraspecific gene flow in *F. virgniana* and *F. chiloensis* including potentially novel evidence of gene flow in the Hawaiian *F. chiloensis* subsp. *sandwicensis* population.

## Materials and Methods

### Plant Material and Genotyping

We collected data from 67 wild octoploid individuals from *Fragaria virginiana* and *Fragaria chiloensis* genotyped with the 50K SNP array developed by Hardigan et al. (2020a) and filtered to remove markers with >5% missing data and markers that were not polymorphic, which brought the final number to 32,200. Detailed sampling and sequencing information can be found in Hardigan et al. (2020a). These samples include four subspecies of *F. virginiana:* subsp. *virginiana* (FVV), *glauca* (FVG), *platypetala* (FVP), and *grayana* (FVY) and four subspecies of *F. chiloensis:* subsp. *chiloensis* (FCC), *pacifica* (FCP), *lucida* (FCL), and *sandwicensis* (FCS). All *F. virginiana* samples are from North America with subsp. *grayana* and subsp. *virginiana* concentrated on the East coast and subsp. *glauca* and subsp. *platypetala* in the West coast. *F. chiloensis* subsp. *pacifica* and *F. chiloensis* subsp. *lucida* are from North America, *F. chiloensis* subsp. *chiloensis* is from South America, and *F. chiloensis* subsp. *sandwicensis* is from Hawaii (Fig 1A, Table 1). See supplemental table 1 for information for detailed geographic locations. A total of seven samples were removed, three (PI 551951, PI 616777, PI616778) were listed in the USDA GRIN database as *F × ananassa* and four (PI 551735, PI 551736, PI 236579, PI616554) were shown to be hybrids in previous analysis (Hardigan et al., 2020b) and USDA GRIN metadata.

**FIGURE 1.**
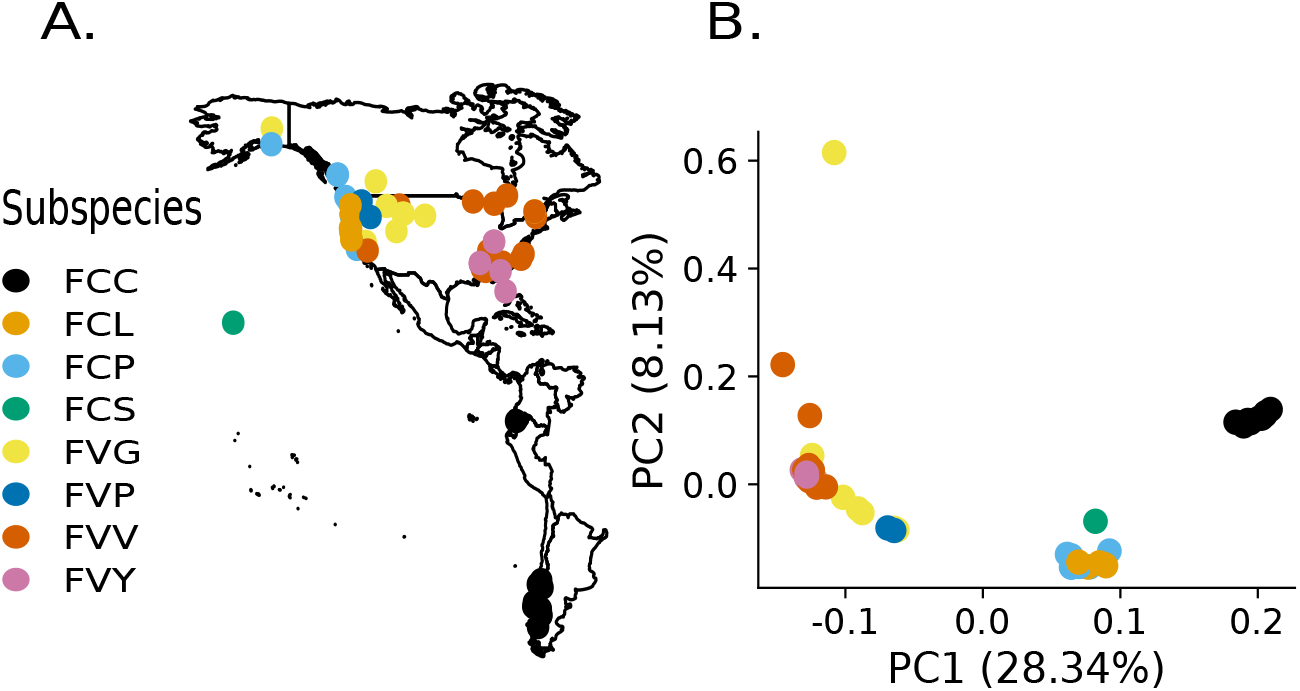
Geography, genetic structure, and phylogenetics of wild octoploid strawberr. **A** Geographic breakdown of sampled wild octoploid strawberry with location data as reported from USDA NPGS GRIN-Global Passport data. For countries without exact latitude and longitude coordinates, coordinates of the described regions were used (Supp table 1) **B** Genetic structure of all wild octoploid samples from PCA 32,200 SNPs.

**TABLE 1.**
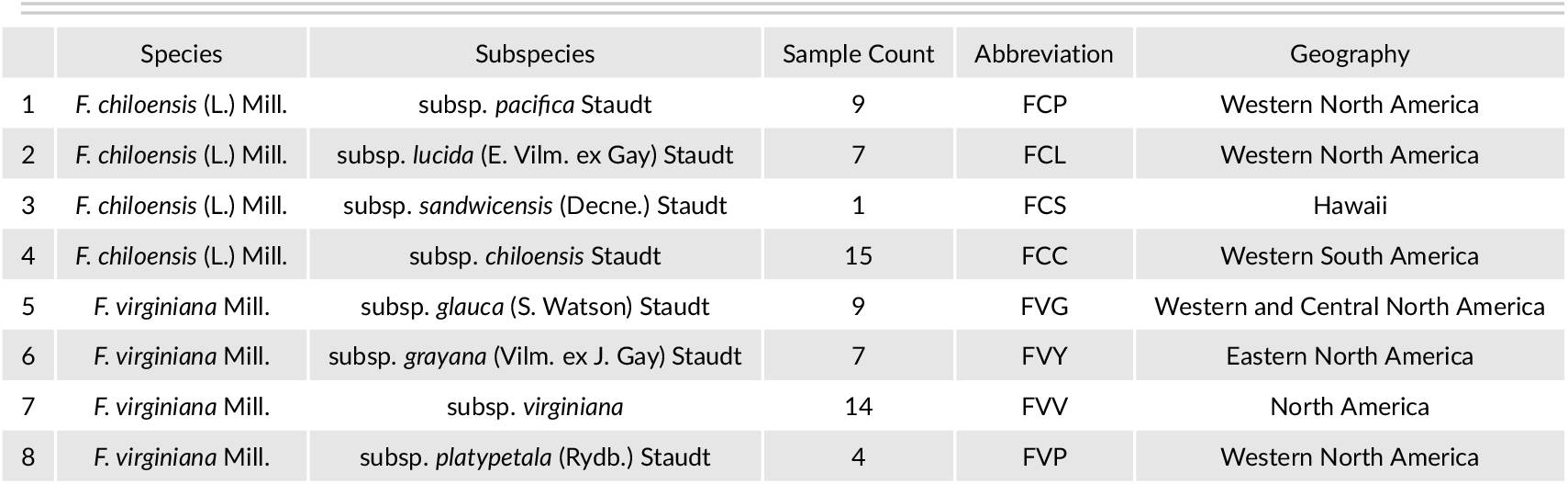
Taxonomy of *Fragaria* octoploids and description the study samples

|  | Species | Subspecies | Sample Count | Abbreviation | Geography |
| --- | --- | --- | --- | --- | --- |
| 1 | *F. chiloensis* (L.) Mill. | subsp. *pacifica* Staudt | 9 | FCP | Western North America |
| 2 | *F. chiloensis* (L.) Mill. | subsp. *lucida* (E. Vilm. ex Gay) Staudt | 7 | FCL | Western North America |
| 3 | *F. chiloensis* (L.) Mill. | subsp. *sandwicensis* (Decne.) Staudt | 1 | FCS | Hawaii |
| 4 | *F. chiloensis* (L.) Mill. | subsp. *chiloensis* Staudt | 15 | FCC | Western South America |
| 5 | *F. virginiana* Mill. | subsp. *glauca* (S. Watson) Staudt | 9 | FVG | Western and Central North America |
| 6 | *F. virginiana* Mill. | subsp. *grayana* (Vilm. ex J. Gay) Staudt | 7 | FVY | Eastern North America |
| 7 | *F. virginiana* Mill. | subsp. *virginiana* | 14 | FVV | North America |
| 8 | *F. virginiana* Mill. | subsp. *platypetala* (Rydb.) Staudt | 4 | FVP | Western North America |

### Phylogenetic Analysis

*Rubus occidentalis* was chosen as the outgroup. We modified the Camarosa v1 octoploid strawberry genome assembly (Edger et al., 2019) to contain ‘N’ characters at SNP locations targeted by 50K array marker probes in order to force SNP calling at those sites against the outgroup genome assembly. We used the ‘nucmer’ function (-maxgap 2500 -minmatch 11 -mincluster 25) in MUMMER v3 (Kurtz et al., 2004) to align the *Rubus occidentalis* v1.1 (outgroup) assembly (Jibran et al., 2018) to the modified Camarosa v1 assembly, and generated SNPs from the alignments using MUMMER’s ‘show-snps’ function to identify the corresponding location of the 50K array SNP sites in the *R. occidentalis* genome sequence. We used BEDTools v2.27 (Quinlan and Hall, 2010) to extract the subset of 50K array SNP sites covered by a single *R. occidentalis* genomic sequence alignment, and then to extract the outgroup nucleotide state at 50K array SNP sites from the *R. occidentalis* v1.1 genome assembly. The *R. occidentalis* nucleotide state at 50K array SNP positions was treated as a homozygous outgroup genotype, except in cases where neither allele measured by the 50K array marker matched the outgroup nucleotide.

We used the coalescent-based Singular Value Decomposition for Quartets method (SVDQuartets) (Chifman and Kubatko, 2014) implemented in PAUP 4.0 (Swofford, 2003) to estimate a phylogeny for the wild octoploid samples and rooted the tree with *Rubus occidentalis*. SVDQuartets computes singular value decomposition (SVD) scores from a matrix of SNP allele frequencies to estimate splits for four taxa trees, called quartets. A species tree is estimated by sampling all combinations of these quartets, inferring a tree for each one, and using an algorithm to combine quartets into a species tree. We evaluated all possible quartets and produced 100 bootstrap replicates. Clades were defined based on recorded subspecies and sample geography.

### Genetic Structure

We generated a genotype matrix from the 32,200 SNPs to resolve the genetic structure of the octoploid *Fragaria* individuals. Population structure was evaluated in two ways. First, we used the R package SNPRelate (Zheng et al., 2012) to perform principal component analysis and plotted the results with ggplot2 (Wickham, 2016). Second, we applied the Bayesian clustering method fastStructure (Raj et al., 2014). We tested K = 2 to 11 clusters with ten cross-validations for each K using the default convergence criterion and prior. The optimal K value was estimated with the chooseK tool contained in the fastStructure package. Results were visualized in Rv 3.6.3 using the conStruct package (Bradburd et al., 2018) and aligned to the phylogenetic tree in Inkscape (Inkscape Project).

### Admixture and Introgression Analysis

We interrogated populations for evidence of admixture and introgression in two ways. We first used the model based approach from TreeMix to identify likely admixture events (Pickrell and Pritchard, 2012). TreeMix infers relationships between populations by modeling genetic drift at genome-wide polymorphisms. It does so by comparing the covariance structure modeled by a computed dendrogram to the observed covariance between populations. If populations are more closely related than the modeled bifurcating tree then an admixture event in the history of those populations is inferred and TreeMix adds a migration edge to the phylogeny. Aspects of the migration edges like position and directions provide further information about the admixture event. For example, if an edge originates from more basal positions on the phylogenetic network it indicates that admixture occurred deeper time or was from a more diverged population. TreeMix was used to create a maximum likelihood phylogeny of the nine subspecies. We rooted the graphs with *F. virginia* subsp. *grayana*, used blocks of 25 SNPs, estimated evolutionary history with one to five migration events, and used the -global option and -se option to calculate standard errors of migration proportions and the -noss option to prevent overcorrection for subspecies with small sample sizes. The OptM R package (Fitak, submitted) was used to determine the optimal number of migration edges. To induce enough variation to assess an optimal model, TreeMix was run with 0-5 migration edges, and block sizes from 200 to 4000 SNPs, in increments of 200 per iteration. The output files from TreeMix were used as input for OptM, and we used the Evanno method to estimate the proportion of variance explained by different number of migration edges. We considered both the point at which 99.8% of variance was explained by the model and the point at which the ad hoc statistics Δm was maximized to assess the optimal number of migration edges. Finally, we ran TreeMix with and without the *F. chiloensis* subsp. *sandwicensis* sample to interrogate its population history as well as broader relationships between the two *Fragaria* species.

For the second strategy, we used several tree-based statistics in ADMIXTURETOOLS2 (https://uqrmaie1.github.io/admixtools/index.html) that screen for excess allelic correlation across branches that do not match a null expectation of a tree-like population history. The first used is the D-statistic, defined as:

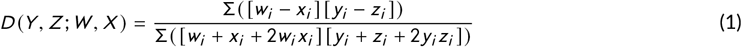

where Y, Z, W, and X are the specific populations, sample frequencies are denoted with lowercase *y, z, w, x* and *i* is an individual locus.

The D-statistic was used in two ways. First we analyzed the full set of reciprocally monophyletic trees of the form ((Y,Z),(W,X)) where *F. chiloensis* subspecies are (Y,Z) and *F. virginiana* are (W,X). Statistical significance was determined based on reported Z-scores, which represent deviations of the D-statistic from zero in units of the standard error. We chose a significance threshold of Z > |3|. Significant deviations from 0 can be interpreted as indicating that (Y,Z) is not a clade relative to (W,X). Positive values indicate excess affinity between Y and W, Z and X, or both and negative values excess affinity between Y and X, Z and W, or both, and therefore rejects reciprocal monophyly.

Second, we applied the D-statistic in a similar fashion to Brandvain et al. (2014) and set up a series of calculations where Y is the *F. chiloensis* subsp. textitpacifica populations which are largely sympatric with *F. virginiana* populations, Z is the *F. chiloensis* subsp. *lucida*, which is more allopatric to *F. virginiana* but is geographically close to *F. chiloensis* subsp. *pacifica*, W is all the *F. virginiana* subspecies, and X is the allopatric South American *F. chiloensis* subsp. *chiloensis*, which is geographically distant from *F. chiloensis* subsp. *pacifica*. This design allowed the testing of gene flow between sympatric *F. chiloensis* and *F. virginiana* populations, indicated by significant positive values of the D-statistic.

Finally, the three population (*f*_3_) test for admixture was performed. The *f*_3_ statistic looks at a three branched phylogeny (A,B;C) and tests whether population C is a mixture of populations A and B. We calculated *f*_3_ statiscs for all populations in cases where either N. American or S. American *F. chiloensis* subspecies were population A and Eastern or Western *F. virgniana* subspsecies were population B. In addition to the *f*_3_ statistic, a Z-score was calculated which represents deviation of the *f*_3_ statistic from zero in units of the standard error. We considered an *f*_3_ statistic as evidence of admixture if the three population test showed a Z-score lower than −3. Importantly, only negative *f*_3_ statistics are unambiguous evidence for admixture.

Additionally *f*_3_ statistics can be used to construct an admixture graph. We took estimated *f*_3_-statistics and the topology of an admixture graph and used the ADMIXTOOLS2 shiny app run_shiny_admixtools() function to find the edge weights that minimize the difference between fitted and estimated *f*_3_-statistics, and summarize that difference in a likelihood score. We considered a good model to be one for which predicted and empirical *f*_3_ and *f*_4_ statistics deviate from 0 by Z-scores < |3| and which have a significantly lower likelihood score than competing graphs. We evaluated whether a likelihood score was significantly different between competing graphs by repeated bootstrap resampling of SNP blocks. We constructed our admixture graph by starting with the topology supported by our phylogenetic analysis, running optimizations to find topologies with lower likelihood scores, comparing fit to an admixture graph with an added admixture event and repeating until modeled *f*_3_ and *f*_4_ statistics fit observed f-statistics with the lowest observed likelihood score. In order to avoid over-fitting with too many model parameters we did not add any more admixture events once all modeled *f*_3_ and *f*_4_ statistics fit the observed f-statistics. If likelihood scores did not significantly differ among graphs with non-significant *f*_3_ and *f*_4_ statistics, preference was given for migrations that matched estimates from Treemix, the three-population test, or both.

## Results

### Genetic Structure and Phylogenetic Relationships

Our combined genetic structure and phylogenetic analyses support three distinct genetic clusters among the 60 *Fragaria* accessions (Fig1B, Fig2A). PCA showed that *F. virgniana* and N. and S. American *F. chiloensis* subspecies formed distinct clusters (Fig 1B). These clusters predominately separated along PC1 which accounted for 28.3% of the variation in the G-matrix.

**FIGURE 2.**
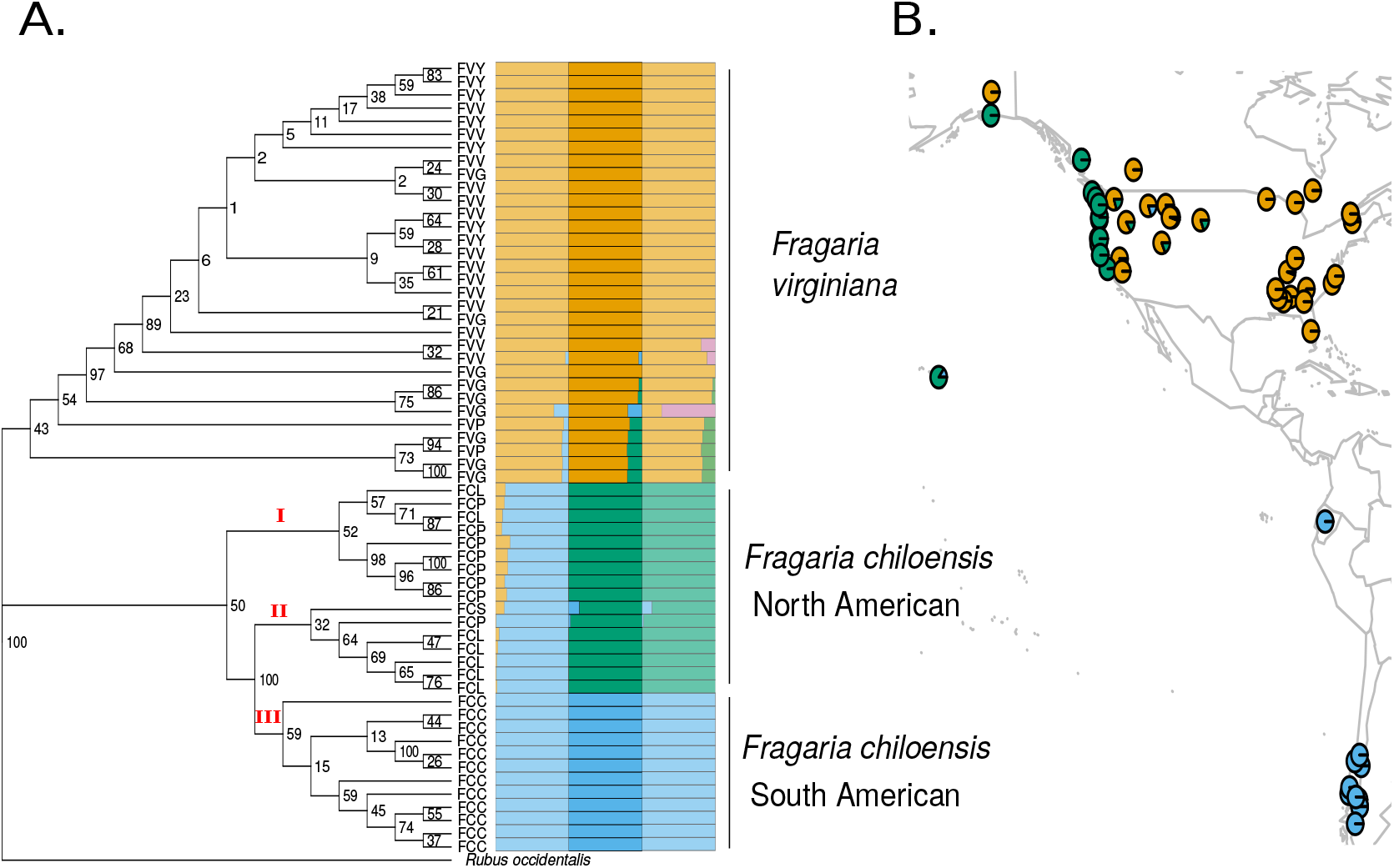
**A.** SVDQuartets phylogeny, excluding samples identified as hybrids in previous analyses, based on 6880 SNPs and using *Rubus occidentalis* as an outgroup, paired with genetic structure estimated from fastSstructure at K=2, 3 and 4. Numerals I, II, and III in red mark clades of S. American *F. chiloensis*. **B.** geographic location of wild octoploid strawberry samples with their inferred structure components at K=3.

From fastStructure using 32,000 SNPs with K=2, we found that *F. virginiana* and *F. chiloensis* were largely separated into distinct clusters, although *F. chiloensis* individuals from North America show admixture with *F. virginiana*, whereas *F. virginiana* individuals from the Pacific Northwest and Canada show admixture with *F. chiloensis* (Fig 2A). At K=3, *F. chiloensis* subspecies were divided into those native to North America (subsp. *pacifica* and *lucida*) and those native to South America (subsp. *chiloensis*). Additionally, the *F. chiloensis* subsp. *sandwicensis* accession from Hawaii was placed with the North American *F. chiloensis* cluster, although with sizable contribution from the South American *F. chiloensis* cluster (Fig 2A). At K=4, three *F. virginiana* individuals show partial contribution from this fourth component. Only one individual showed > 50% contribution from the fourth component. It’s unclear whether this fourth component is a distinct subpopulation or attributable to introgression from other *Fragaria* species. While the ChooseK script from FastStructure indicated that K=4 maximizes the marginal likelihood and best explains the structure of the data, individuals did not have > 80% contribution from the components added at K=4, so K=3 was chosen as the best representative of the data.

In order to incorporate *Rubus occidentalis* as an outgroup genotype in our analysis of strawberry markers, we used whole-genome alignment to identify *R. occidentalis* nucleotide states in sequences corresponding to the 50K array SNP sites we assayed in the octoploid strawberry genome. We used MUMMER to perform whole-genome alignment of the *R. occidentalis* genome to the four octoploid strawberry subgenomes. We then identified the subset of SNP positions in the octoploid genome that are targeted by probes on the 50K SNP array and that were covered by a single *R. occidentalis* alignment from an ancestrally related chromosome based on a previous analysis of chromosome synteny by Hardigan et al. (2020a). The nucleotide state at the position in the *R. occidentalis* genome assembly corresponding to 50K array marker SNP sites was assigned as a homozygous outgroup genotype for the corresponding markers, except in cases where the R. occidentalis allele did not match one of the two nucleotide states assayed by the marker probe. Indel sites were ignored. In total 9,840 of the filtered, polymorphic markers were assigned a corresponding *R. occidentalis* outgroup genotype, of which 6,687 remained following exclusion of markers with 5% missing data.

Phylogenetic analysis using the 6,687 SNPs shared among *Fragaria* species and *Rubus occidentalis* showed two major clades, with *F. virginiana* sister to all *F. chiloensis* (Fig2A). The bootstrap support within the *F. virginiana* clade was frequently low (24/31 <75%), and all subspecies appear paraphyletic; however, a well supported branch separated the Eastern N. American subsp. *virginiana* and subsp. *grayana* individuals from individuals of the Western N. American subspecies (subsp. *glauca* and subsp. *platypetala*). Within the *F. chiloensis* clade there are three subclades, marked on the phylogeny (Fig 2A). Clade I on the phylogeny is primarily comprised of subsp. *pacifica* individuals from the coast of Alaska and the Pacific Northwest of the US and Canada)Fig 2A,B) and is sister the remaining *F. chiloensis* populations. Notably, the branch separating Clade I from Clades II and III has low bootstrap supporter (50%) suggesting the splitting of these clades is better represented as a polytomy. Clade II is predominately comprised of subsp. *lucida* and *sandwicensis* individuals from coastal California (Fig 2A,B) and is sister to all of the S. American *F. chiloensis* with strong bootstrap support (100%). Finally, clade III is a well resolved South American *F. chiloensis* clade. These results suggest that North American *F. chiloensis* subspecies are paraphyletic, while the S. American subspecies, subsp. *chiloensis* is monophyletic.

### Admixture and Introgression Analysis

We next ran TreeMix with the eight representative subspecies, adding migration edges to the phylogenetic tree until model fit was optimized. We found that the TreeMix model with two migration edges maximized the likelihood of the model and was the point at which 99.8% variance was explained. We found strong evidence of gene flow from a common ancestor of the *F. chiloensis* clade into the Hawaiian *Fragaria chiloensis* subsp. *sandwicensis*. The second migrations estimated by TreeMix suggests admixture between a population ancestral to contemporary *F. chiloensis* subsp. *chiloensis* and the N. American *F. chiloensis* subsp. *pacifica* (Fig 2A). The migrations after these were from Eastern N. American *F. virginiana* to *F. virginiana* subsp. *glauca*, from an ancestral *F. chiloensis* subsp. *chiloensis* into *F. chiloensis* subsp. *lucida* and from an ancestral *F. chiloensis* subsp. *pacifica* into *F. virginiana* subsp. *virginiana* (Supp Fig 1).

Because there is only one individual from *F. chiloensis* subsp. *sandwicensis* sampled and the signal for admixture was strong we ran TreeMix a second time excluding this sample to get an estimates of broader relationships between the two *Fragaria* species. The optimal number of migrations in this analysis was two based on the ad hoc Δm metric and the percent of variance explained by the model. The first is from the internal branch between the Western and Eastern *F. virgniana* subspecies, representing a population ancestral to the Western *F. virginiana* populations, into *F. chiloensis* subsp. *pacifica* (Fig 3B). Interestingly, the tree topology in this case shows *F. chiloensis* subsp. *pacifica*, rather than subsp. *lucida*, as sister to *F. chiloensis* subsp. *chiloensis*. The second migration was a population ancestral to the Eastern *F. virgniana* subsp. *virgniana* populations into the Western *F. virgniana* subsp. *glauca*. Subsequent migration events were similar to those inferred from the previous analysis including *F. chiloensis* subsp. *sandwicensis* (Supp Fig 2)

**FIGURE 3.**
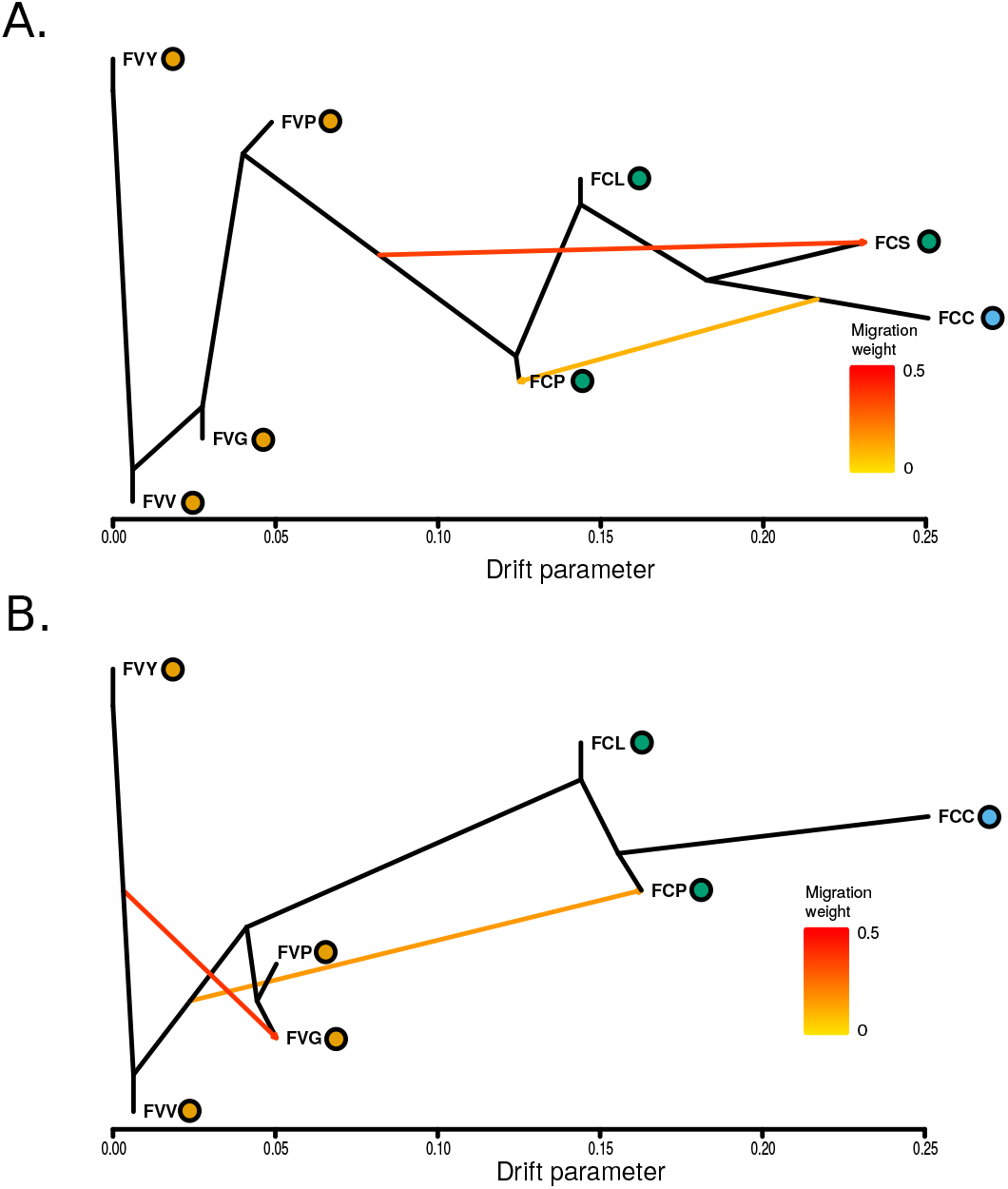
TreeMix analysis with optimal number of migrations including (**A.**) and excluding (**B.**) *F. chiloensis* subsp. *sandwicensis*. Colored dots indicate population membership assigned by FastStructure.

In addition to the model based approach of TreeMix, we used analyses that employed three and four branched population trees for signals of gene flow. Our first implementation D-statistic allowed us to infer whether a given four branch tree follows a topology of reciprocal monophyly ((Y,Z);(W,X)). Both positive and negative significant D statistics results with Z-scores > |3| were taken as rejection of the null hypothesis that (Y,Z) forms a clade relative to (W,X). Positive values suggests gene flow between Y and W, or Z and X and negative values suggest gene flow between Y and X or Z and W. Out of the 18 total combinations of *F. chiloensis* and *F. virginiana* four population trees, four had Z-scores > |3|, indicating they reject a simple tree structure of reciprocal monophyly (Table 2). These involved trees with N. and S. American *F. chiloensis* subspecies for W and X, and Eastern and Western *F. virginiana* subspecies for Y and Z. All four significant results had negative Z-scores, in these cases suggesting gene flow between either N. American *F. chiloensis* and Eastern *F. virgniana* or between S. American *F. chiloensis* and Western *F. virgniana*.

**TABLE 2.** D-statistic test for reciprocal monophyly of *F. chiloensis* and *F. virgniana*

|  | W | X | Y | Z | D | SE | Zscore | Significant? | n |
| --- | --- | --- | --- | --- | --- | --- | --- | --- | --- |
| 1 | FCC | FCP | FVV | FVG | -0.0402 | 0.00878 | -4.585 | Yes | 25165 |
| 2 | FCC | FCP | FVY | FVG | -0.0415 | 0.00971 | -4.275 | Yes | 25165 |
| 3 | FCC | FCL | FVY | FVG | -0.0319 | 0.00946 | -3.372 | Yes | 25165 |
| 4 | FCC | FCL | FVV | FVG | -0.0301 | 0.00911 | -3.306 | Yes | 25165 |
| 5 | FCP | FCL | FVV | FVG | 0.0135 | 0.00489 | 2.756 | No | 25165 |
| 6 | FCC | FCP | FVP | FVY | 0.0321 | 0.01182 | 2.714 | No | 25120 |
| 7 | FCC | FCP | FVP | FVV | 0.0307 | 0.01131 | 2.713 | No | 25120 |
| 8 | FCP | FCL | FVY | FVG | 0.013 | 0.00539 | 2.411 | No | 25165 |
| 9 | FCP | FCL | FVP | FVV | -0.0184 | 0.00873 | -2.106 | No | 25120 |
| 10 | FCP | FCL | FVP | FVY | -0.0179 | 0.00894 | -2 | No | 25120 |
| 11 | FCC | FCL | FVP | FVG | -0.0147 | 0.01083 | -1.354 | No | 25120 |
| 12 | FCC | FCL | FVP | FVY | 0.0173 | 0.01285 | 1.344 | No | 25120 |
| 13 | FCC | FCL | FVP | FVV | 0.0153 | 0.01259 | 1.215 | No | 25120 |
| 14 | FCP | FCL | FVP | FVG | -0.0063 | 0.00671 | -0.933 | No | 25120 |
| 15 | FCC | FCP | FVP | FVG | -0.0083 | 0.01036 | -0.798 | No | 25120 |
| 16 | FCC | FCL | FVY | FVV | -0.0022 | 0.00629 | -0.347 | No | 25165 |
| 17 | FCC | FCP | FVY | FVV | -0.0017 | 0.00576 | -0.303 | No | 25165 |
| 18 | FCP | FCL | FVY | FVV | -4e-04 | 0.00459 | -0.08 | No | 25165 |

Note: Z-scores represent deviations from 0 in terms of standard error. D-statistics with Z scores > |3| are considered significant and a rejection of reciprocal monophyly of *F. chiloensis* and *F. virginiana*.

As an additional test for admixture, we used the D-test structure from Brandvain et al. (2014) to test specifically for gene flow between *F. chiloensis* subsp. *pacifica* and *F. virginiana* subspecies. These tested trees all showed significantly positive Z-scores (>10) suggesting gene flow between all *F. virginiana* subspecies and *F. chiloensis* subsp. *pacifica* (Table 3).

**TABLE 3.**
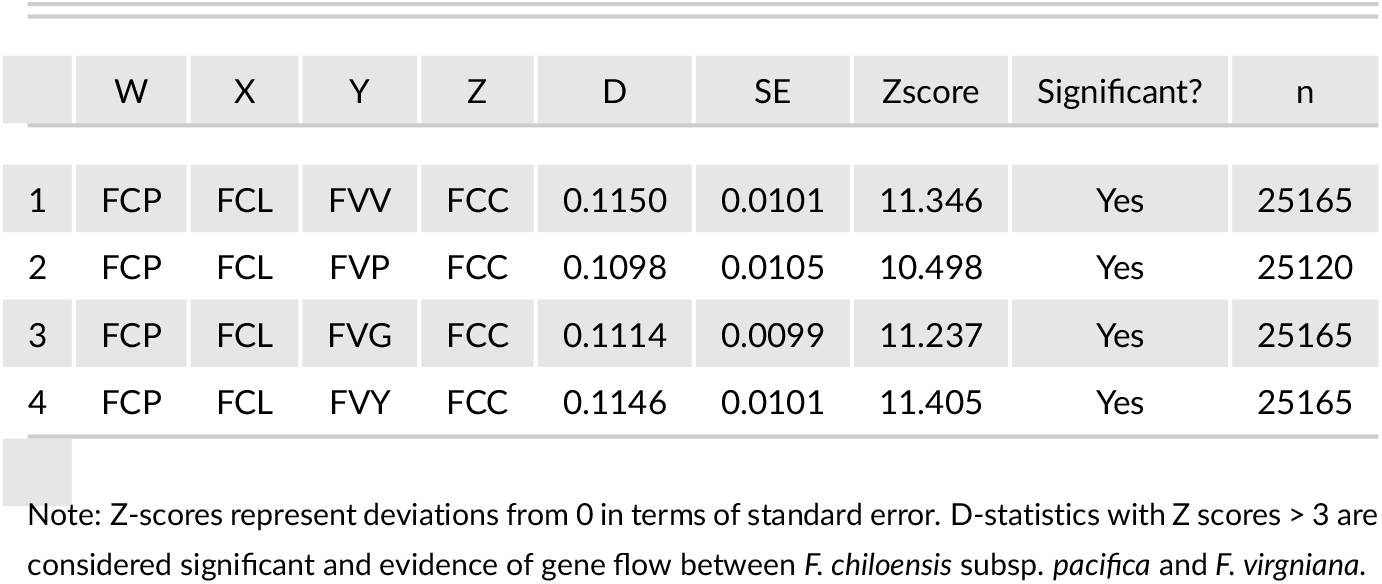
D-statistic Test for gene flow between *F. chiloensis subsp. pacifica* and *F. virginiana*

*f*_3_ statistics from the three-population test provide additional evidence for admixture. Based on trees showing Z-scores less than −3, *F. chiloensis* subsp. *pacifica* and *F. virginiana* subsp. *glauca* are suggested to be admixed (Fig 4). *F. chiloensis* subsp. *pacifica* showed admixture from both Eastern and Western *F. virgniana* populations but only when paired with S. American *F. chiloensis*. Meanwhile, *F. virginiana* subsp. *glauca* showed evidence of admixture from all 3 *F. chiloensis* subspecies, but only when paired with Eastern *F. virgniana* populations. Although there was large negative *f*_3_ values for *F. chiloensis* subsp. *lucida*, the deviation was not significantly different from 0.

**FIGURE 4.**
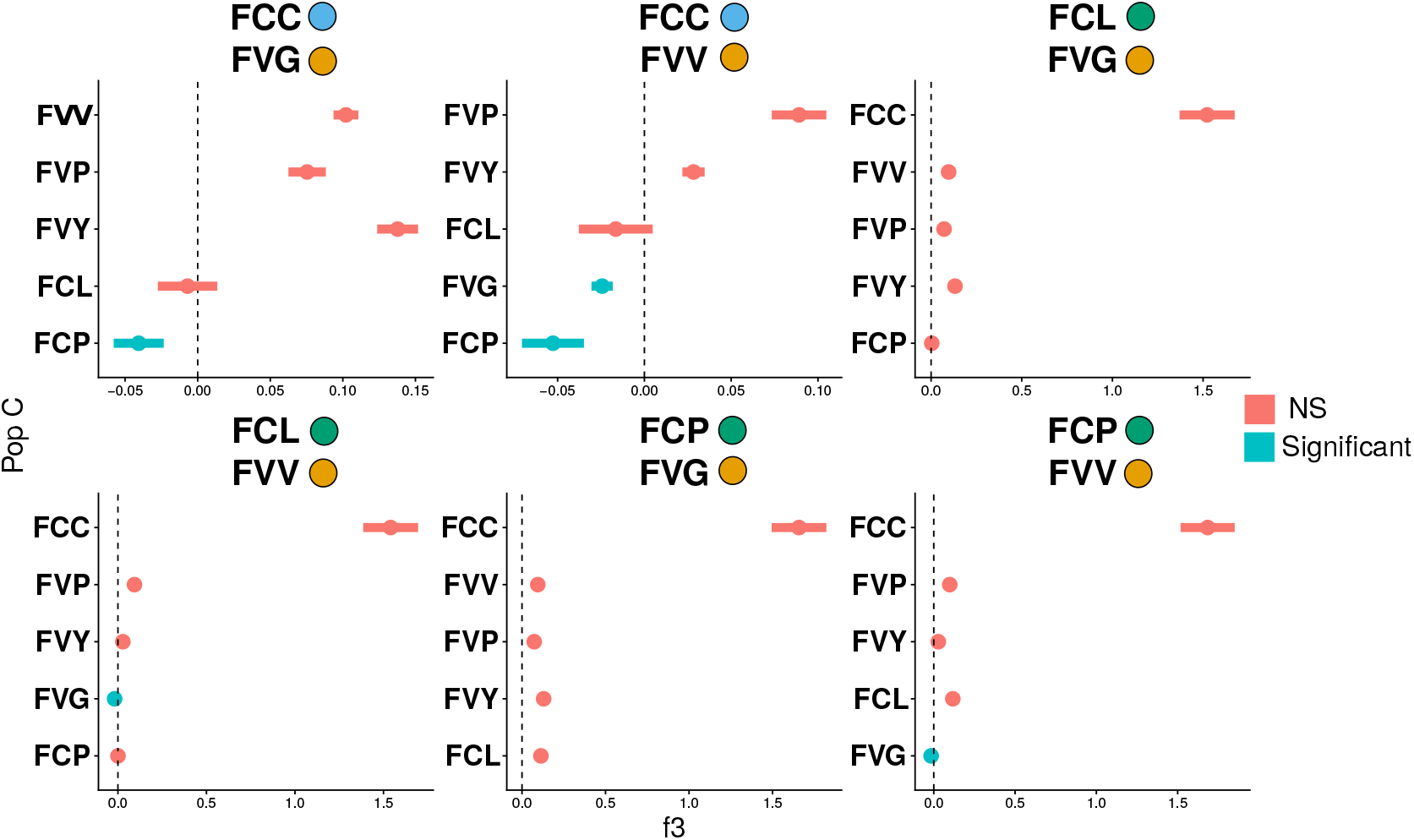
Three Population Test (*f*3) statistics. *f*3 statistics are shown for all subspecies (population C) with S. American *F. chiloensis* subsp. *chiloensis* or N. American *F. chiloensis* subsp. *pacifica* and subsp. *luicida* as population A and Eastern *F. virgniniana* subsp. *virginiana* and Western *F. virginiana* subsp. *glauca* as population B. Colored dots indicate population membership assigned by FastStructure. Points represent mean *f*3 statistics and lines the standard error. Only *f*3 statistics with Z-scores less than −3 are considered statistically significant and are marked in blue.

**FIGURE 5.**
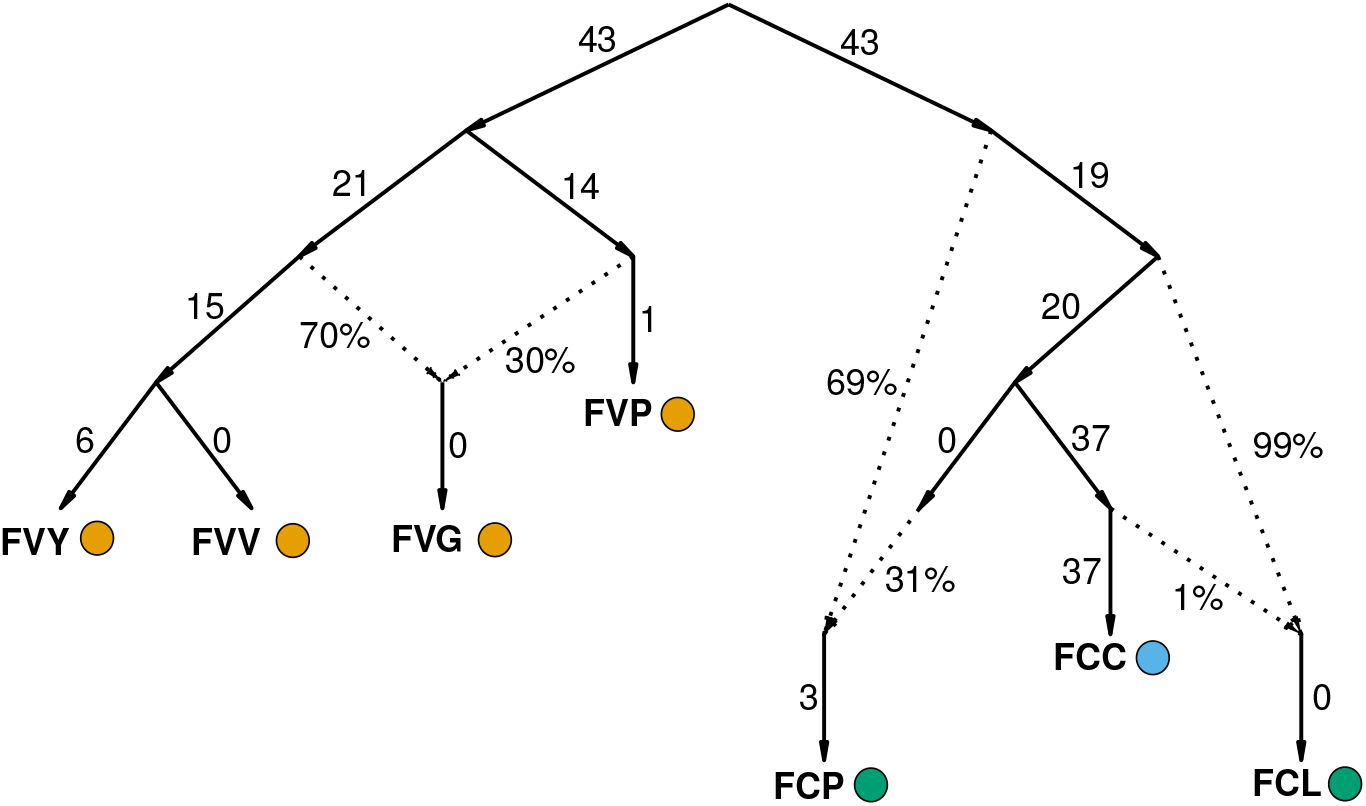
Admixture Graph. Best fitting admixture graph showing the three main admixture events between *Fragaria* species and subspecies. Admixture events are marked by the dotted arrows. Numbers proximal to solid lines are drift lengths of branches and percentages proximal to dotted lines are admixture weights.

Finally, these *f*3 trees were combined to estimate a full admixture graph. Three admixture events significantly improved fit over a base tree and provided a statistical fit to the observed *f*3 and *f*4 statistics from the data (Fig 4). These events mirror the events inferred from TreeMix providing several lines of convergent evidence in support of these admixture events. The three events include *F. virginiana* subsp. *glauca* having contributions from an ancestral branch of the Eastern *F. virgniana* clade and a population ancestral to *F. virginiana* subsp. *platypetala*. Next, *F. chiloensis* subsp. *pacifica* is estimated to have having contributions from an ancestor of the S. American *F. chiloensis* subsp. *chiloensis* subspecies and a population ancestral to all *F. chiloensis* populations. Finally, *F. chiloensis* subsp. *lucida* is estimated having contributions from a population ancestral to *F. chiloensis* subsp. *chiloensis* and a population ancestral to the divergence of *F. chiloensis* subsp. *chiloensis* and *F. chiloensis* subsp. *lucida*.

## Discussion

Wild octoploid *Fragaria* are complex species with a rich history of range expansions, speciation, and hybridization that has been difficult to dissect. Recent advances in genomic resources have allowed a more detailed investigation of the relationships between the two wild octoploid species that are progenitors to the agriculturally important *F*. × *ananassa*.

### Origins and dispersal of *Fragaria* octoploids

The SVDQuartets phylogeny provides the first in depth look at the evolutionary relationships of wild octoploids using nuclear DNA. Based on this tree, both *F. virginiana* and *F. chiloensis* are monophyletic sister species with strong bootstrap support. These results strongly suggest both species diversified after colonizing new ecological niches across the Western hemisphere..

These results diverge from a previous analysis using plastid DNA (Dillenberger et al., 2018) which did not find monophyly for *F. virginiana* and found *F. chiloensis* to be sister to *F. virginiana* subsp. *platypetala*. The difference in these results may be due to the lack of other *Fragaria* species sampled in this study which would have made the *F. virginiana* clade paraphyletic, as was observed with the inclusion of octoploid *F. × ananassa* subsp. *cuneifolia* and the decaploid *F. cascadensis* in Dillenberger et al. (2018). The lack of monophyly with plastid DNA might also be due to hybrdization and introgression that is thought to be common between *Fragaria* species. However, it is unlikely that sampling alone changed the relationship of *F. virginiana* subsp. *platypetala* as a common ancestor to all *F. chiloensis* so the differences in the previous results may largely be due to hybridization and gene flow obscuring the species relationship inferred from plastid DNA.

Contrary to the species level, only one of eight subspecies, *F. chiloensis* subsp. *chiloensis* was monophyletic. However there was some signal of geographic patterning in the phylogeny and genetic structure analysis. South American *F. chiloensis* subspecies were monophyletic and were identified as a distinct genetic cluster at K=3 using structure. North American *F. chiloensis* were also identified as a distinct genetic cluster, however phylogenetically they formed two clades, one of predominately *F. chiloensis* subsp. *pacifica* which is found along the Alaskan, Canadian, and Pacific North West coast, and one of predominately *F. chiloensis* subsp. *lucida* which is found along the West coast of the United States from the Pacific North West to Southern California.

Notably, the *F. chiloensis* clades reflect the geographic expansion of the species, with the most Northern populations (*F. chiloensis* subsp. *pacifica*) in clade I sister to clades II and II which are both more southern populations, and the Pacific North West/California Coast populations in clade II sister to the S. American populations in clade III. These results suggest that there was a gradual range expansion and diversification of the species as it moved South. This is bolstered by the observation from Hardigan et al. (2020b) showing almost complete overlap of S. American alleles and N. American *F. chiloensis* alleles, where all S. American alleles are a subset of N. American alleles. The reduced heterozygosity and longer LD decay of South American *F. chiloensis* compared to North American *F. chiloensis* observed by Hardigan et al. (2020b) is further evidence of this kind of range expansion.

All of these signals are consistent with a single origin and bottleneck from range expansion. Additionally the one sample from Hawaii was within clade II, the Coastal California clade, with notable genetic contribution from the S. American *F. chiloensis* clade. These two results are consistent with *F. chiloensis* in South American and Hawaii likely being from independent population movements (Hancock and Prince, 2020). The results here would suggest these two populations were dispersed independently from Southern California coastal populations related to modern day *F. chiloensis* subsp. *lucida*.

### Intra- and interspecific admixture among *Fragaria* populations

The *Fragaria* genus is notorious for interspecific hybridization and polyploidization, with several allopolyploids ranging from tetraploid to decaploid, and the main crop garden strawberry being the result of spontaneous hybridization of *F. virginiana* and *F. chiloensis*. However, characterization of hybridization and gene flow in the wild among these octoploids has not been heavily investigated using genomic methods until now. Results from distinct analyses to detect gene flow converge on inter- and intra-specific gene flow among *Fragaria* populations. Four populations in particular show evidence of being admixed.

First, multiple lines of evidence support admixture for *F. chiloenesis* subsp. *pacifica*. The independent methods converge on contributions from S. American *F. chiloensis* subspecies and an ancestral populations with more recent common ancestry with *F. virginiana* populations. In both implementations of TreeMix, *F. chiloensis* subsp. *pacifica* had ancestry contributions from *F. chiloensis* subsp. *chiloensis* and an ancestral *F. virginiana* population. Similarly the three-population test for admixture and the D-statistic test that investigated admixture between *F. chiloensis* subsp. *pacifica* and *F. virginiana* show evidence of admixture from all *F. virginiana* subspecies. The admixture with deeper ancestral populations likely explains why *F. chiloensis* subsp *pacifica* showed significant D-statistics and *f*3 statistics from all *F. virginiana* populations and it is likely these results are either from admixture deep in the *F. virgniana* phylogeny or a population very deep in the *F. chiloensis* clade when genetic variation from *F. virginiana* was still present. These models are more parsimonious than multiple admixture events in each subspecies and are more concordant with TreeMix and the admixture graph. Although there have not been any time-calibrated phylogenies that can accurately date the diversification of *Fragaria* octoploids, these results potentially indicate that *F. chiloensis* subspecies may have diverged earlier than *F. virginiana* subspecies. Additionally, based on where these migration edges occurred, it is likely this admixture occurred prior to the *F. chiloensis* subsp. *chiloensis* movement to S. America, which would suggest the *F. chiloensis* populations began diverging prior to the substantial geographic separation currently observed. Alternatively, movement between N. and S. American populations may have been more common at some point in the past.

The next population suggested to be admixed is the Hawaiian *F. chiloensis* subsp. *sandwicensis*. TreeMix inferred admixture from an ancestral *F. chiloensis* population, potentially prior to any geographic or subspecies divergence. This may be due to the region being originally colonized by an older gene pool and having gene flow from more recent populations on the mainland. The genetic structure patterns, and inferred admixture suggests that these Hawaiian populations may be a novel gene pool not fully represented by N. or S. American populations. However, it’s important to note that analyses here are limited in that only one individual from this population is included and only one method was employed to infer admixture events. Although TreeMix is somewhat robust to low sample sizes it will be crucial to increase sampling of this population to improve the resolution of results and further our understanding of its evolutionary history.

All methods employed also converged on *F. virgniana* subsp. *glauca* being admixed. The three-population test methods, admixture graph, and the optimal migrations for the second run of TreeMix excluding the Hawaiian poulation identified admixture among *F. virginiana* subspecies, particular the Eastern and Western populations. Additionally, although gene flow between *F. chiloensis* populations were the only migrations selected as optimal by TreeMix when the Hawaiian *F. chiloensis* subsp. *sandwicensis* was included, this admixture event was inferred by subsequent added admixture edges, providing additional support for this admixture event. These admixture events are estimated to have occurred in the common ancestor of the Eastern *F. virginiana* populations and the other Western populations. *F. virgniana* subsp. *glauca* in particular is known to have a range that spans the West and East coast of the US, overlapping with the other *F. virgniana* populations. This range likely explains why there is consistent gene flow between the East and West coasts. Interestingly, significant D-statistic and three-population test *f*_3_ statistic suggest that *F. virgniana* subsp. *glauca* is also admixed by all *F. chiloensis* subspecies. The admixture graph also shows an admixture edge from deep within the *F. virgniana* clade. This signal may be due to gene flow form an ancestral *F. chiloensis* population, or from an ancestral *F. virginiana* population with genetic variation shared with *F. chiloensis* still segregating.

Our results provides preliminary evidence for admixture in *F. chiloensis* subsp. *lucida*. In the admixture graph, the addition of an admixture event from *F. chiloensis* subsp. *chiloensis* into subsp. *lucida* was necessary for the model *f*_3_ and *f*_4_ statistics to fit the observed data. In both implemtentations of TreeMix, admixture from *F. chiloensis* subsp. *chiloensis* into subsp. *lucida* was inferred, but these events were after the model fit was optimized and so have weaker support. Likewise, the three-population test showed a large negative *f*_3_ statistic when *F. chiloensis* subsp. *chiloensis* was included, but the signal was not significant. These results are suggestive of admixture, but more mixed than other signals and may require followup with expanded sampling of subsp. *lucida*.

The extent of intraspecific gene flow inferred by these methods likely partially explains the paraphyly observed in the phylogeny, although given the robustness of SVDQuartets to gene flow (Long and Kubatko, 2018) and the more recent divergence of these populations, incomplete lineage sorting, subspecies misidentification, or a combination may be more likely. Thoroughly investigating incomplete lineage sorting among *Fragaria* species and subspecies is a promising subject for future phylogenetic studies in this system.

Finally, several lines of converging evidence suggest there has been prominent interspecific gene flow, especially between N. American *F. chiloensis* and *F. virginiana* populations. First, the FastStructure plot identifies signals of N. American *F. chiloensis* ancestry in several Western *F. virgniana* samples and at K=2 N. American *F. chiloensis* from clade I show contribution from the *F. virgniana* population cluster. Additionally, the second application of D-statistics, specifically investigating gene flow between sympatric *F. chiloensis* and *F. virginiana* also found significant signals of gene flow, specifically between *F. chiloensis* subsp. *pacifica* and all *F. virginiana* populations. These results were bolstered by three-population test *f*_3_ statistics. Notably, subsp. *pacifica* show evidence of admixture between South American *F. chiloensis* and various *F. virginiana* subspecies, reflecting patterns of admixture observed when FastStructure was run at K=2 and the admixture inferred from S. American *F. chiloensis* into *F. chiloensis* subsp. *pacifica*. Likewise, *F. virginiana* subsp. *glauca* showed signs of being admixed, with contribution from all *F. chiloensis* subspecies and all other *F. virginiana* subspecies populations. The mixture with deeper ancestral populations likely explains that *F. chiloensis* subsp *pacifica* showed significant *f*3 statistics from all *F. virginiana* populations and *F. virginiana* subsp. *glauca* showed significant *f*3 statistics from all *F. chiloensis* subspecies, as these ancient populations likely shared many alleles with the diverging sister species. While natural hybrids between the *F. chiloensis* and *F. virginiana* have been documented where their ranges overlap in the Pacific Northwest (Hancock Jr and Bringhurst, 1979; Staudt, 1999), these results indicate that there was sustained gene flow between these populations that left a mark on the genome of these species. Additionally the lack of monophyly in the plastid phylogeny of Dillenberger et al. (2018) may be explained by the observed gene flow between these overlapping populations.

### Another genomic tool in our box

Beyond the empirical results presented here, this study also highlights the utility of well designed genotyping arrays for limited phylogenetic and population genomic analyses. With complex genomes, especially one like octoploid *Fragaria* where there are 4 distinct subgenomes, SNP calling and genotyping from reduced representation libraries like RADseq and Genotyping-By-Sequencing contain missing data and can be prone to errors (Blischak et al., 2018), and the ability to distinguish subgenomes equally across the genome may be compromised. Whole-genome resequencing can resolve this better, but is more expensive and may be computationally challenging with complex polyploid genomes. The genotyping array used in this study was designed to capture diversity across subgenomes and identify subgenome-specific SNPs. Nicely designed SNP arrays results in even coverage across all subgenomes with minimal missing data for a relatively small cost (Hardigan et al., 2020a). There are some concerns for their application in population genomic studies. Traditionally genotyping arrays are designed for association studies and will have ascertainment biases that complicate the use of any methods reliant on the site-frequency spectrum (SFS) (Clark et al., 2005). However, many methods, like those used in this study such as tree-based statistics and TreeMix, are robust to many ascertainment schemes (Patterson et al., 2012; Pickrell and Pritchard, 2012) and will allow for cursory examinations of evolutionary relationships between populations in species previously inaccessible to population genomic investigation. Additionally, designing genotyping arrays with population genomic studies in mind may allow for directly addressing ascertainment bias, allowing the use of SFS-based methods like scans for selection or demographic modelling. Reduced representation libraries, whole-genome resequencing, and genotyping arrays have their respective strengths and weaknesses, but recognizing the utility of well designed genotyping arrays in complex polyploid systems may help facilitate future population genomic work.

## Conclusion

Recent developments in genomic resources in *Fragaria* allowed for unprecedented investigation into the origins and evolution of the wild *Fragaria* octoploids. These results helped clarify the phylogenetic relationship of these octoploids, providing strong support they are monophlytic sister species and characterizing the extensive admixture within and among species. In particular, regions of sympatry are marked by extensive interspecific gene flow for both *F. chiloensis* subsp. *pacifica* and *F. virgniana* subsp. *glauca*. We also identified additional cases of admixture for followup studies. The Hawaiian population *F. chiloensis* subsp. *sandwicensis* appears to be a gene pool with admixture from an ancestral *F. chiloensis* population that is worthy of future investigation, and a potential signal of admixture in *F. chiloensis* subsp. *lucida* was found that would both benefit from additional sampling of these populations. This study also highlights the role that inexpensive genotyping arrays and carefully selected analyses can play in evolutionary studies of organisms with large or complex genomes.

## Acknowledgements

This research was supported by grants to KAB from the National Science Foundation (NSF-GRFP DGE-1424871), to PPE from the National Science Foundation (NSF-PGRP #2029959) and United States Department of Agriculture (AFRI #2020-67013-30870), and to SJKfrom the United Stated Department of Agriculture (http://dx.doi.org/10.13039/100000199) National Institute of Food and Agriculture (NIFA) Specialty Crops Research Initiative (#2017-51181-26833), California Strawberry Commission (http://dx.doi.org/10.13039/100006760), and the University of California, Davis (http://dx.doi.org/10.13039/100007707)

## Supporting Information

**Supp. Fig. 1:** TreeMix results modeling 0-5 migration edges, including *F. chiloensis* subsp. *sandwicensis*

**Supp. Fig. 2:** Treemix results modeling 0-5 migration edges, excluding *F. chiloensis* subsp. *sandwicensis*

