## Supplementary figures and images for "Diversification, Spread, and Admixture of Octoploid Strawberry in the Western Hemisphere"

### Supp. Fig. 1

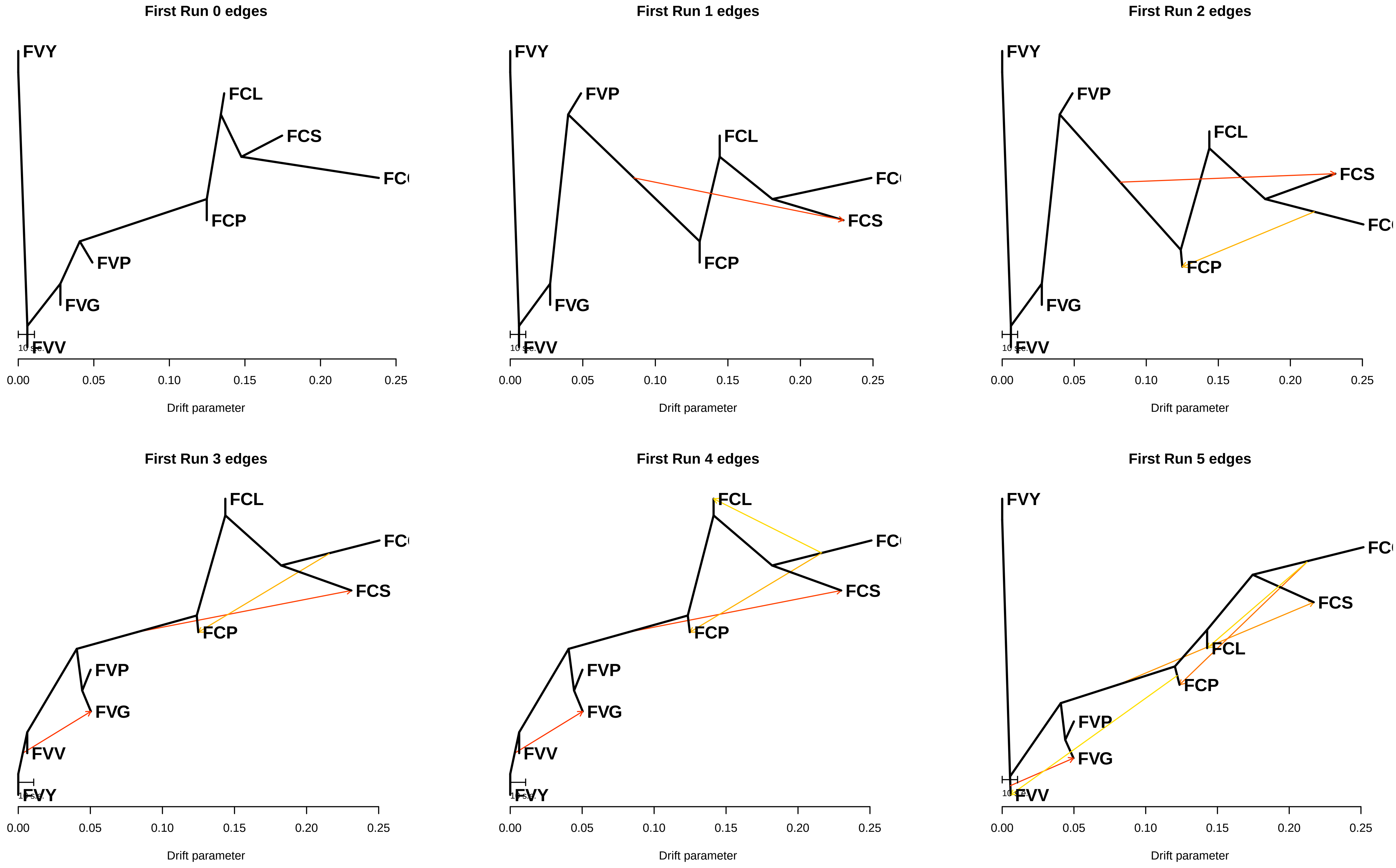

### Supp. Fig. 2

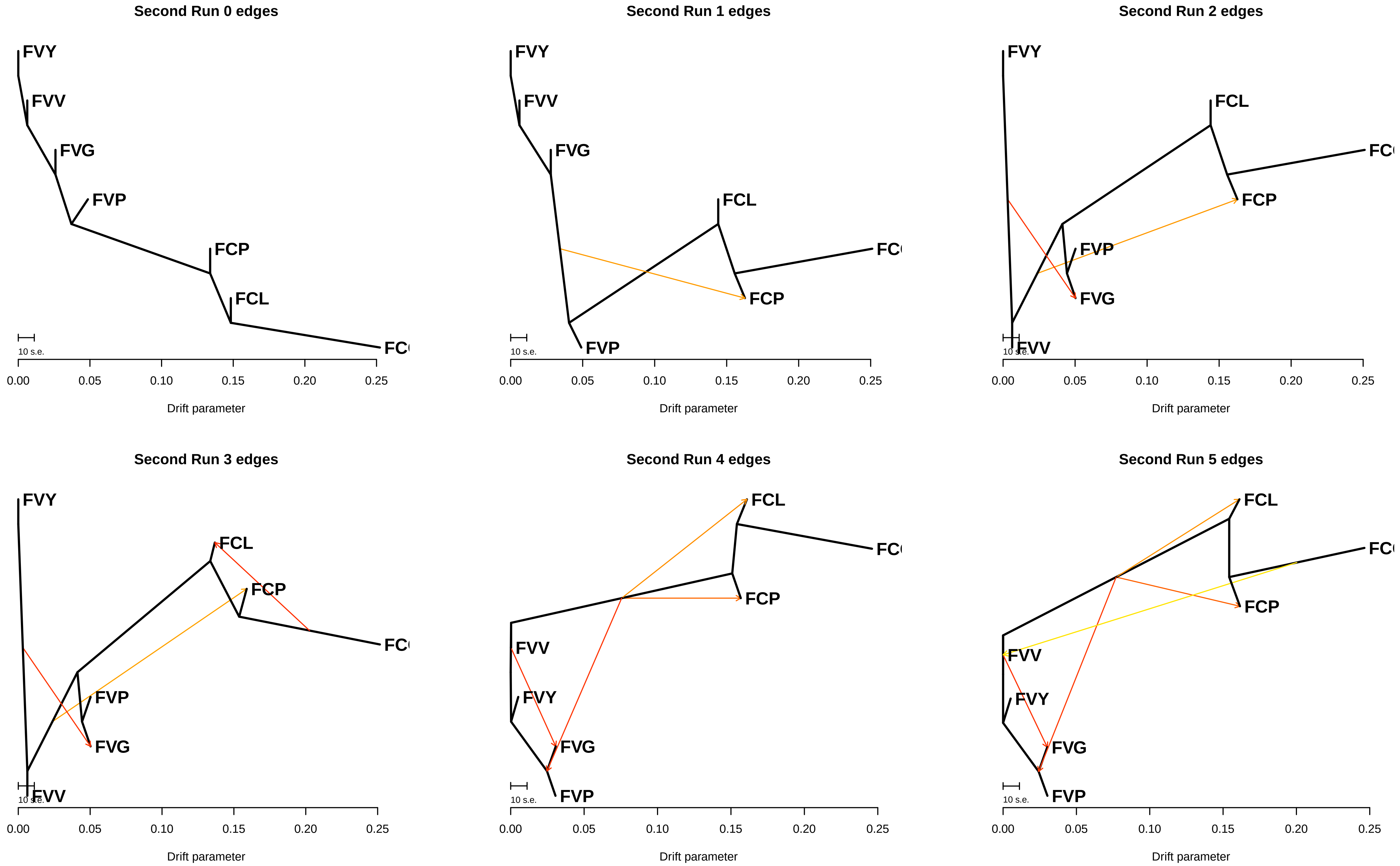
